# Computational Designing of a Multi-Epitope Vaccine Against *Streptococcus pyogenes*

**DOI:** 10.64898/2026.08.02.742269

**Authors:** Sana Hassan, Saleha Mohammed Razaulla, Rajan Kumar Pandey

## Abstract

Group A *Streptococcus (GAS)*, or *Streptococcus pyogenes* is almost exclusive and greatly adapted human pathogen. It causes a wide array of clinical symptoms, ranging from minor infections of the skin and soft tissues to pharyngitis, meningitis, pneumonia, bacteraemia, cellulitis, puerperal sepsis, and necrotising fasciitis. The risk of *S. pyogenes* infection is known to be influenced by several host characteristics, including age, underlying diseases like diabetes, varicella, or skin lesions, both chronic and acute, and certain risk behaviours such as use of drugs. Household size and overcrowding are two environmental factors that significantly affect the transmission of *S. pyogenes*. The majority of cases occur spontaneously in the community, and preventative opportunities are still limited. A large portion of GAS-related mortality is found in low-income areas and communities. Based on aforementioned public health risk, the creation of effective therapeutic vaccines would be an excellent addition to current control measures. The purpose of this work is to address the need for new instruments to aid in the elimination of *S. pyogenes* infections. The discovery of high antigenic regions in several highly conserved proteins brings us one step closer to developing peptide vaccines capable of influencing the different phases of *S. pyogenes* infection, providing more effective defence and greater serotype coverage. This study used various techniques of immunoinformatics to design an effective multi-epitope vaccine that produced neutralising antibodies against multiple strains of *S. pyogenes*.

## INTRODUCTION

Group A Streptococcus (GAS), also known as *Streptococcus pyogenes,* is estimated to currently infect about 18 million humans globally, with a yearly death rate of around 517,000 due to severe manifestations of the resulting diseases (1). The signs and symptoms range from simple skin and soft tissue infections to pharyngitis and more serious conditions like meningitis, pneumonia, bacteraemia, cellulitis, puerperal sepsis, and necrotising fasciitis. The occurrence of streptococcal toxic shock syndrome (STSS) in affected people considerably raises their risk of death (2). Major improvements have occurred in the global epidemiology of GAS and the reporting of its suppurative and non-suppurative infections since the early 1980s (3). The prevalence of invasive *S. pyogenes* infection varies with time and geography, which is likely a reflection of a population’s susceptibility to specific strains as well as the intrinsic diversity in the most prevalent varieties (4). Numerous virulence pathways are involved in the organism’s complex pathogenicity (5). Age, medical conditions like diabetes, varicella, or other acute or chronic skin lesions, or risky behaviours like injecting drugs, are all known to have an impact on the likelihood of contracting *S. pyogenes* infection (6–8). Household size and overcrowding are two environmental factors that are known to significantly affect the transmission of *S. pyogenes* (9). There are still limited chances for prevention and the majority of cases occur spontaneously within a community. Low-income neighbourhoods and locations account for a significant amount of Group A Streptococcus-related mortality (10). Infections with *S. pyogenes* and their after-effects are more common in socioeconomically disadvantaged communities. Although *S. pyogenes* infection can affect people of any age, they are more common in children aged between 5 to 15 years. It is also known to be one of the highest contributors to deaths due to bacterial sepsis. Pharyngeal infections have been described as a “hazard” for school-age children and are more prevalent in those older than three (11). The relatively low incidence of infection in newborns may indicate a protective, transplacental acquired immunity.

*S. pyogenes* cocci are round to ovoid, gram-positive, non-motile bacteria having M1, M3, M5, M28, M49, and M89 serotypes (12). The *S. pyogenes* M1 strain has 1,852,442-bp genome sequence that includes 1,752 predicted protein-encoding genes. Of them, nearly a third have no obvious function, while the remaining belong to established categories of known microbial activity. Other genes that encode proteins associated with acute glomerulonephritis or rheumatic fever have been identified. These genes are most likely involved in the “molecular mimicry” of host traits by microorganisms(13). *S. pyogenes* strains are divided into two classes based on the post-infectious sequelae they are associated with: class I, which have an immune determinant on the C-repeat conserved region of the M protein. The class is known to cause rheumatic fever after streptococcal infection. Class II causes acute glomerulonephritis and lacks the mentioned immune determinant on the surface-exposed M protein (13). Since it contributes to the virulence of *S. pyogenes*, the M protein has been a major target in vaccine research since the 1940s. Conventional methods, which demonstrated notable effectiveness in human studies, concentrated on the entire M protein, were then narrowed down to the polypeptides of the N-terminal along with regions of C-repeat peptides. As the *emm* types and the resulting N- terminals of the M protein show a high degree of variation, vaccines developed to target that region use a multivalent strategy, as in the case of StreptAnova (14). Despite several promising routes, the pipeline for *S. pyogenes* vaccines is currently quite lacking, particularly in comparison with strategies for other infectious illnesses with a substantial global disease burden (15). Using *in-silico* techniques of immunoinformatics will aid in the rapid designing of a potential multi-epitope vaccine. This approach is known to be more economical and successful than the traditional laboratory procedure (16, 17), although the labour-intensive process of choosing the best subunits from a large number of predictions is one of the issues facing modern vaccine development. Despite their value, these techniques are unable to immediately forecast or choose the best vaccine candidates, which frequently results in excessive computational work.

This study began with utilization of various immunoinformatic tools to predict potential B-and T-cells epitopes followed by designing subunit vaccine. The overall stability and capacity to elicit a robust immunological response was determined using various tools for the subunit vaccine. Although we got promising results, including high antigenicity and immunogenicity profiles, evaluating this subunit vaccine using in vitro and in vivo studies will be required to validate its protective efficacy and safety.

## METHODOLOGY AND MATERIALS

### Sorting proteins for vaccine designing

The complete amino acid sequences of the five *S. pyogenes* proteins emm6, emm5, emm12, emm24, and emm3 were chosen as part of the M Protein. The gene sequences were then obtained in FASTA format from the NCBI Protein Database (https://www.ncbi.nlm.nih.gov/protein). The International Nucleotide Sequence Database Collaboration’s nucleotide and protein sequence databases include organism names and classifications for each sequence, according to the National Centre for Biotechnology Information’s (NCBI) classification (19).

### Predictive analytics for conservancy evaluation

The Clustal Omega is a multiple-sequence alignment tool that creates alignments between three or more biological sequences which can be either nucleic acids or proteins (20). It uses the mBED algorithm to calculate guide trees, which allows it to handle many tens of thousands of DNA/RNA or protein sequences. Clustal Omega was used to align all sorted *S. pyogenes* sequences. Those peptides located in regions with strong conservation values were selected for the vaccine designing. The Clustal Omega server can be accessed at https://www.ebi.ac.uk/jdispatcher/msa/clustalo.

### Predicting linear B-cell epitopes

B-cell epitopes are the antigenic segments, particularly the solvent-exposed regions, that are recognized by B-cells and then bound by immunoglobulins. These epitopes are crucial for developing vaccines with high immunogenicity through antibody production (21). The linear B-cell epitopes were predicted using the online ABCpred tool (22). ABCpred server uses an Artificial Neural Network (ANN) to predict the B cell epitope or epitopes in an antigen sequence (https://webs.iiitd.edu.in/raghava/abcpred/ABC_submission.html). ABCPred classifies epitopes and non-epitopes using a recurrent neural network to improve accuracy (22). It is very important to have computational techniques that can accurately anticipate linear B-cell epitopes in protein sequences, as humoral immune response involving antibodies is dependent on the antigenic epitopes and their interaction with B-cell receptors (23).

### Prediction of cytotoxic T lymphocytes (CTL) epitopes

In infected cells, pathogenic peptide epitopes are processed and presented to the CTLs by MHC class I molecules. This binding of peptides to HLA-1 is a highly selective process, which is why effective vaccine design requires accurate estimates of cytotoxic T lymphocyte (CTL) epitopes (24). The CTL epitopes from specific proteins were predicted using NetCTL-1.2 (https://services.healthtech.dtu.dk/services/NetCTL-1.2/), a high-sensitivity technique based on artificial neural networks (25). Two supertypes of HLA, A3 and B7, were selected for epitope identification of CTL epitope prediction, as they show wide population coverage despite the high polymorphism of HLA alleles (26). The CTL epitopes were chosen using the combined score and half-maximal inhibitory concentration (IC50) <50 nm(27). According to the criteria, high scores indicated good binding, and an IC50 of less than 50 nm suggested the epitope with the highest receptor affinity. The values selected indicate how likely a peptide is to bind to the MHC Class I molecule and be recognised by the CTLs (CD8+ T cells).

### Prediction of helper T-cell (HTL) epitopes

Helper T Lymphocytes (HTL) influence the activity of various cells of the immune system and use their T-cell receptor (TCR) to recognize epitopes presented by specific MHC class II molecules(28). The predictions of possible HTL epitopes were achieved using the NetMHCII 2.2 Server (http://www.cbs.dtu.dk/services/NetMHCII/). This server can predict peptides having good binding affinity to an MHC II molecule by using ANNs. The database is calibrated on a large amount of data that includes more than 500,000 values from binding affinity (BA) and eluted ligand mass spectrometry (EL). The service also uses ANNs to predict peptide binding to the HLA-DR, HLA-DQ, and HLA-DP alleles (29). The HTL epitopes were selected using the IC50 value of <50 nm and the lowest percentile ranks. Low affinity is indicated by an IC50 value <5000 nM, and intermediate affinity is indicated by a value <500 nM (30). Therefore, the percentile rank may directly correlate with the IC 50 and inversely correlate with the epitope’s affinity.

### Predicting epitopes that induce IFN-**γ**

Important for both innate and adaptive immune responses, the cytokine interferon-gamma (IFN-γ) stimulates macrophages and natural killer cells and improves their response to MHC antigens. Apart from its broad-spectrum antiviral effect, IFN-γ also boosts MHC’s responsiveness to antigens. Interferon-gamma-inducing epitopes can be predicted and designed using the IFNepitope server (http://crdd.osdd.net/raghava/ifnepitope/scan.php), which uses a support vector machine (SVM) (31). This server creates overlapping sequences. It also uses information that includes an MHC class-II binder, which can induce IFN-γ and activate T-helper cells. After the selected HTL epitopes were uploaded to the IFNepitope server, the IFN-γ inducing epitopes were exclusively chosen for developing the final vaccine.

### Construction of multi-epitope vaccine candidate sequence

The vaccine candidate sequence was created using a combination of high-affinity HTL-, CTL-, and B-cell epitopes. To increase the vaccine’s immunogenicity, an adjuvant, Beta-defensin (Uniprot ID P81534), was added to the vaccine sequence (32). The Uniprot server can be accessed at http://www.uniprot.org/. Additionally, a KK linker (33) was added between the adjuvant and B-cell epitope, and intra-B-cell epitopes, an AAY linker (34) between CTL epitopes, and GPGPG linkers (35) were used between HTL epitopes.

### Prediction of the vaccine protein antigenicity

Antigenic reactivity, also referred to as antigenicity, is the capacity of a peptide to be recognized by complementary immune receptors or antibodies at their specific binding sites (36). Antigenicity prediction allows the peptide’s antigenicity to be enhanced by the structure-based rational design concept by increasing the degree of steric complementarity between it and a single monoclonal antibody. To predict the antigenicity of the identified proteins, the tool VaxiJen v2.0 was utilized, which was the first alignment-independent protective antigen prediction. It was reported that the technique has a prediction accuracy of 70% to 89%. The server can be used independently or in conjunction with prediction techniques based on alignment at ( http://www.jenner.ac.uk/VaxiJen). VaxiJen v2.0 is based on the auto-cross-covariance (ACC) conversion of protein sequences into uniform vectors that depict the primary properties of amino acids (37).

### Prediction of the vaccine protein allergenicity

It is essential to assess an antigen’s allergenicity before it is administered to humans. Numerous bioinformatics tools and databases are available for this purpose, taking into consideration a protein’s potential for allergens and non-allergens (38). The first alignment-free server for *in-silico* allergen prediction based on the primary physicochemical characteristics of proteins is called AllerTOP. It is an allergy prediction server that uses machine learning and is openly accessible. It was reported that the prediction accuracy of this instrument was 85.3% (39). The degree of allergenicity of the vaccination was evaluated by the AllerTOP v.2.0 and AllergenFP 1.0 web servers. The web server for AllerTOP v.2.0 can be accessed at (https://www.ddg-pharmfac.net/AllerTOP/data.html). In addition to allergenicity, AllerTOP can predict whether an allergen will be exposed by food, inhalation, or toxicity (40).

### Prediction of the vaccine protein toxicity

CSM-Toxin was utilized to predict the toxicity of vaccine peptides. CSM-Toxin is a unique *in-silico* protein toxicity classifier that depends entirely primary sequence of the given protein. The CSM-Toxin effectively identified the potential toxicity of proteins and peptides, with an MCC of up to 0.66 across cross-validation and several non-redundant blind tests (41). The software can be accessed at https://biosig.lab.uq.edu.au/csm_toxin/.

### Physicochemical properties of the vaccine protein

Comparative analysis of the physicochemical characteristics of a protein is important to understand its function and its molecular evolution. The physicochemical characteristics of a single protein sequence can be determined using a variety of offline and online techniques. The web server ProtParam (http://web.expasy.org/protparam/) was used to evaluate the physicochemical parameters (42). ProtParam is a tool that enables the calculation of different physical and chemical properties for a protein sequence or a specific protein found in UniProtKB. The molecular weight, theoretical pI, extinction coefficient, predicted half-life, atomic and amino acid composition instability index, aliphatic index, and grand average of hydropathicity (GRAVY) are among the calculated parameters through the ProtParam server.

### Solubility prediction of the vaccine peptide

In both industrial and medicinal applications, protein solubility is an important characteristic. Despite a growing understanding of the relevant physicochemical aspects, prediction remains difficult. The Protein-Sol web server was used for the prediction of the solubility of proteins. It can be accessed at (http://protein-sol.manchester.ac.uk) (43).

### Prediction of secondary structures

The PSIPRED protein structure prediction server was utilized to evaluate the secondary structures of the vaccine construct (44). PSIPRED also offers a variety of precise protein annotation tools that facilitate the completion of highly scalable biological investigations. With it, users can submit a protein sequence, select a prediction, and receive the results both graphically over the web or email. The server can be accessed at (http://bioinf.cs.ucl.ac.uk/psipred/).

### Prediction, improvement, and validation of tertiary structure

Proteins’ three-dimensional structures can be accurately predicted from their amino acid sequences (45). It has been demonstrated that DeepMind’s cutting-edge machine learning model, AlphaFold, is highly effective for this purpose (46). The server can be accessed at https://deepmind.google/technologies/alphafold/ and was utilized to predict the protein’s structure. The Galaxy Web server was utilized to refine the three-dimensional model obtained for the multi-epitope vaccination peptide (https://galaxy.seoklab.org/cgi-bin/submit.cgi?type=REFINE). The refinement technique employed by the GalaxyRefine web server has been effectively validated in CASP10. This process involves rebuilding side chains, repacking them, and using molecular dynamics modelling to relax the overall structure. The CASP10 evaluation revealed that this approach performed best in enhancing the quality of the local structure (47). The Ramachandran plot was generated by the MolProbity software https://molprobity.biochem.duke.edu, which allowed us to validate the structure(48). Based on the side chain’s van der Waals radius, the Ramachandran plot illustrates the energetically permitted and disallowed dihedral angles of an amino acid, psi (ψ) and phi (φ). The quality of the modelled structure is determined by the percentage of residues in allowed and restricted regions, which are reported in the AlphaFold results

### Molecular dynamic simulations

Molecular dynamics simulations were performed using GROMACS 2021.5(49), to investigate the refined binding mode, stability, and binding affinity between the TLR4 and the vaccine construct. The signalling pathway of TL4 plays an essential role in the activation of the innate immune system. The simulations employed the AMBER ff99sb force field parameters, similar to an earlier study(50). The systems were solvated in a cubic simulation box with a 10 Å buffer using the TIP3P water model, and counter ions were added to neutralise the system. Energy minimisation was done using the steepest descent algorithm for 5000 steps, followed by the conjugate gradient method for 2000 steps.

To equilibrate the systems, 1 ns of NVT (constant number of particles, volume, and temperature) and 1 ns of NPT (constant number of particles, pressure, and temperature) simulations were conducted. The equilibrated system was then subjected to a 100 ns production molecular dynamics simulation. A cut-off distance of 1.0 nm was applied for short-range interactions, while long-range electrostatic interactions were computed using the particle mesh Ewald (PME) method (51) with a Fourier spacing of 0.16 nm and an interpolation order of 4. The simulation trajectories and system snapshots were analysed and visualized using the built-in GROMACS gmx tools (52).

The AMBER suite’s gmx_MMPBSA v1.6.360. provided the MMPBSA.py script used to assess the binding affinity between the TLR and the vaccine construct, through the calculation of binding energy (53). The final equilibrated portion of the MD simulation trajectory, corresponding to the last 50 ns, was used for these calculations. Entropy contributions were excluded from this analysis due to their high computational cost.

### Predictive analytics of immune simulations

Immunogenicity refers to a vaccine’s ability to induce an immunological response, particularly the development of antibodies that identify and neutralize the target pathogen. *In-silico* immunological simulations were employed to further describe the immunogenicity and immune response of the produced multi-epitope vaccine construct. Both parameters were analysed using the C-ImmSim server, which is a position-specific scoring matrix (PSSM) (54). It predicts immunological interactions using position-specific score matrices for peptide prediction based on machine learning methods. The C-ImmSim model defines the humoral and cellular barriers of the mammalian immune system against vaccine production. The C-ImmSim server can be found at (http://www.cbs.dtu.dk/services/C-ImmSim-10.1/).

### Codon Optimization and Reverse Translation

In order to ensure an efficient expression of the designed vaccine in E.coli, the selected expression vector, reverse translation, and codon optimization were conducted with the Java Codon Adaptation Tool (JCat) server (http://www.prodoric.de/JCat). Reverse translation translates the protein sequence to the corresponding DNA strand. Codon optimization aids in adapting the chosen sequence to align with the host’s preferred usage of codons, thus increasing overall expression and avoiding undesired transcription termination (55).

## RESULTS

### Protein sequence collection and primary analysis

To create a potential multi-epitope vaccination against *S. pyogenes*, the amino acid sequences for five proteins were extracted from the NCBI Database. Proteins with sequences shorter than 100 amino acids were eliminated to identify epitopes for vaccine development. The sequences were divided into two groups to ensure broad vaccine coverage. Group 1, contained the most common and epidemiologically similar *emm* types, widely associated with throat infection. Group 2, comprised of *emm* types that are less common and are part of a single protein *emm* cluster. Proteins emm6 (SPP08089), emm12 (TRQ54508), and emm3 (TRQ05464) were part of Group 1, while Group 2 consisted of emm5 (SPP02977) and emm24 (SPP12379).

### Predictive analytics for conservancy evaluation

Many sequence analysis techniques are based on multiple sequence alignments. The progressive alignment heuristic is used to calculate the majority of alignments. Clustal Omega is a technique that produces precise alignments and can swiftly align nearly any number of protein sequences. The multiple sequence alignment was completed using the Clustal Omega tool. In the first step, the sequences of emm6, emm12, and emm3 from group 1 were run on the software. The sequence common to all group 1 proteins was extracted. To confirm that the selected protein sequence matches exactly in all three of group 1’s proteins, another round of multiple sequence alignment was conducted. The results obtained were accurate, and the sequence (SEASRKGLRRDLDASREAKKQVEKDLANLTAELDKVKEEKQISDASRQGLRRDLD ASREAKKQVEKALEEANSKLAALEKLNKELEESKKLTEKEKAELQAKLEAEAKALK) was chosen. In step two, the multiple sequence alignment was performed on the group 2 proteins, emm5 and emm24. The sequence common to all group 2 proteins was extracted. To confirm that the selected protein sequence matches exactly in both of group 2’s proteins, another round of multiple sequence alignment was conducted. The results obtained were accurate, and the sequence (LRRDLDASREAKKQLEAEHQKLEEQNKISEASRQSLRRDLDASREAKKQLEAEHQK LEEQNKISEASRQSLRRDLDASREAKKQVEKALEEANSKLAALEKLNKELEESKKLT EKEKAELQAKLEAEAKALKEKLAKQAEELAKLRAGKASDSQTPDAKPGNKAVPGK GQAPQAGTKPNQNKAPMKETKRQLPSTGETANPFFTAAALTVMATAGVAAVVKRK EEN) was chosen.

### Predicting linear B-cell epitopes

The ABCpred software was utilized to choose linear B-cell epitopes with variable residue lengths. A window length of 16 mer and a threshold of 0.8 were selected for prediction. The B-cell epitopes were chosen based on the highest scores, indicating a greater likelihood that the peptide is a linear B-cell epitope. Identification and characterization are essential for the development of vaccines, immunodiagnostic procedures, and antibody manufacturing. For the five selected proteins, the ABCpred server predicted 117 B-cell epitopes. The 14 epitopes with scores higher than 0.8 were accepted, and subsequently tested for antigenicity using VaxiJen., Table 1 shows the details of all selected B-cell epitopes.

**Table 1:**
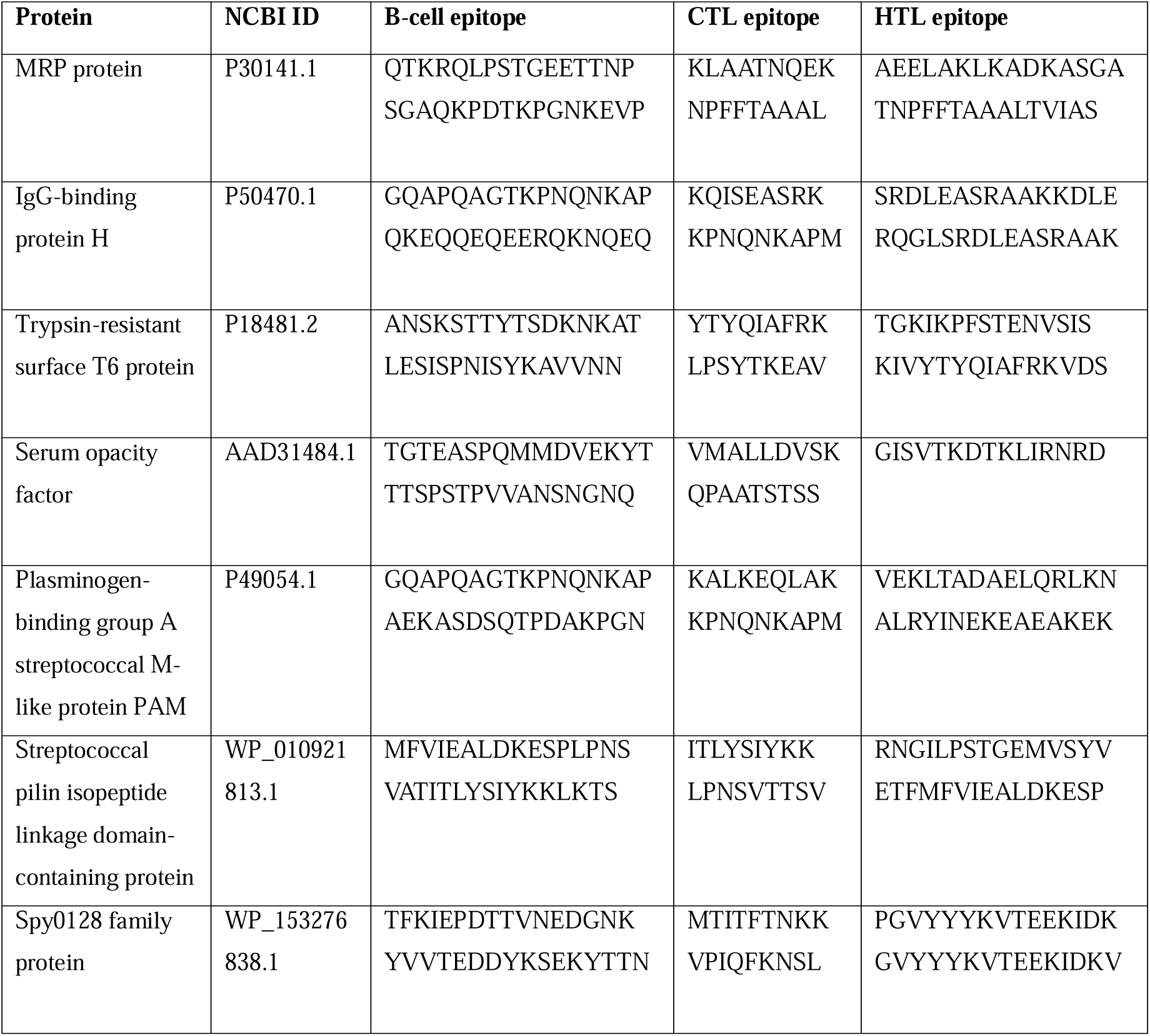
Predicted linear B-cell and T-cell epitopes selected for designing the peptide vaccine against *S. pyogenes*. The order of the epitopes corresponds to their positions in the final chimera design.

### Prediction of cytotoxic T lymphocytes (CTL) epitopes

With the default threshold score of 0.750000 for epitope identification, the NetCTL 1.2 server predicted 107 CTL (9-mer) ligands for the five chosen proteins. 14 CTL-epitopes were selected due to their high binding affinity or immunogenicity scores, which indicated a good IC50 of less than 50 nm. Additionally, the A3 and B7 supertype CTL values were chosen. Table 1 shows the details of all selected CTL-cell epitopes.

### Prediction of helper T-Cell (HTL) epitopes

Prediction of HTL epitopes was done using the NetMHCII 2.2 web server based on their IC50 scores. For this, five alleles of HLA were selected, with the peptide length of the epitopes fixed at 15 mers. A total of 13 high-binding HTL epitopes were chosen due to their low IC50 scores, indicating a strong binding affinity to the MHC Class II molecules. Table 1 displays the details of all selected HTL epitopes. The chosen epitopes were identified as high-binding MHC-II epitopes for the human alleles HLA-DR, HLA-DQ, and HLA-DP.

### Predicting epitopes that induce IFN-**γ**

The IFN epitope server uses the MERCI and SVM software for prediction of epitopes. All 13 HTL epitopes chosen for the primary vaccine sequence were found to have the ability to induce IFN**-**γ based on their positive scores.

### Construction of multi-epitope vaccine candidate sequence

The chimeric vaccine was designed using 14 linear B-cell epitopes, 14 CTL epitopes, and 13 HTL epitopes. The selected epitopes are listed in order as presented in Table 1. KK, GPGPG, and AAY linkers were used to fuse the predicted peptide sequences comprised of linear B-cell, CTL, and HTL epitopes, respectively. The Beta-defensin of Homo sapiens (Human) (Uniprot id: P81534) was chosen as an adjuvant, which is a 50S ribosomal protein that was attached to the N-terminal of the vaccine construct using an EAAAK linker (56) to enhance antigen-specific immune responses. It was selected as the adjuvant due to its small size and non-toxic nature. It also acts as a natural immunostimulant and has properties that activate the Toll-like receptor (TLR), in addition to improving vaccine immunogenicity by enhancing antibody titre, cell proliferation, and cytokine release. The EAAAK linker was used for adjuvant separation as it creates a rigid helical structure and minimizes interactions between the epitopes. The KK linkers were used between the B-cell epitopes to maintain the flexibility and solubility of the short B-cell epitopes. AAY linkers were placed between the CTL epitopes, as they generate natural CTL epitopes that can be effectively presented by MHC class I molecules. Lastly, we added the GPGPG linkers between the HTL epitopes to facilitate proper processing and presentation by the MHC class II molecules. In order to ease the identification and purification of the protein, a 6xHis tag was added to the C-terminal. The final vaccination peptide was composed of 751 amino acid residues taken from 41 merged peptide sequences. The sequence is represented in Figures 1 (A) and 1 (B).

**Figure 1.**
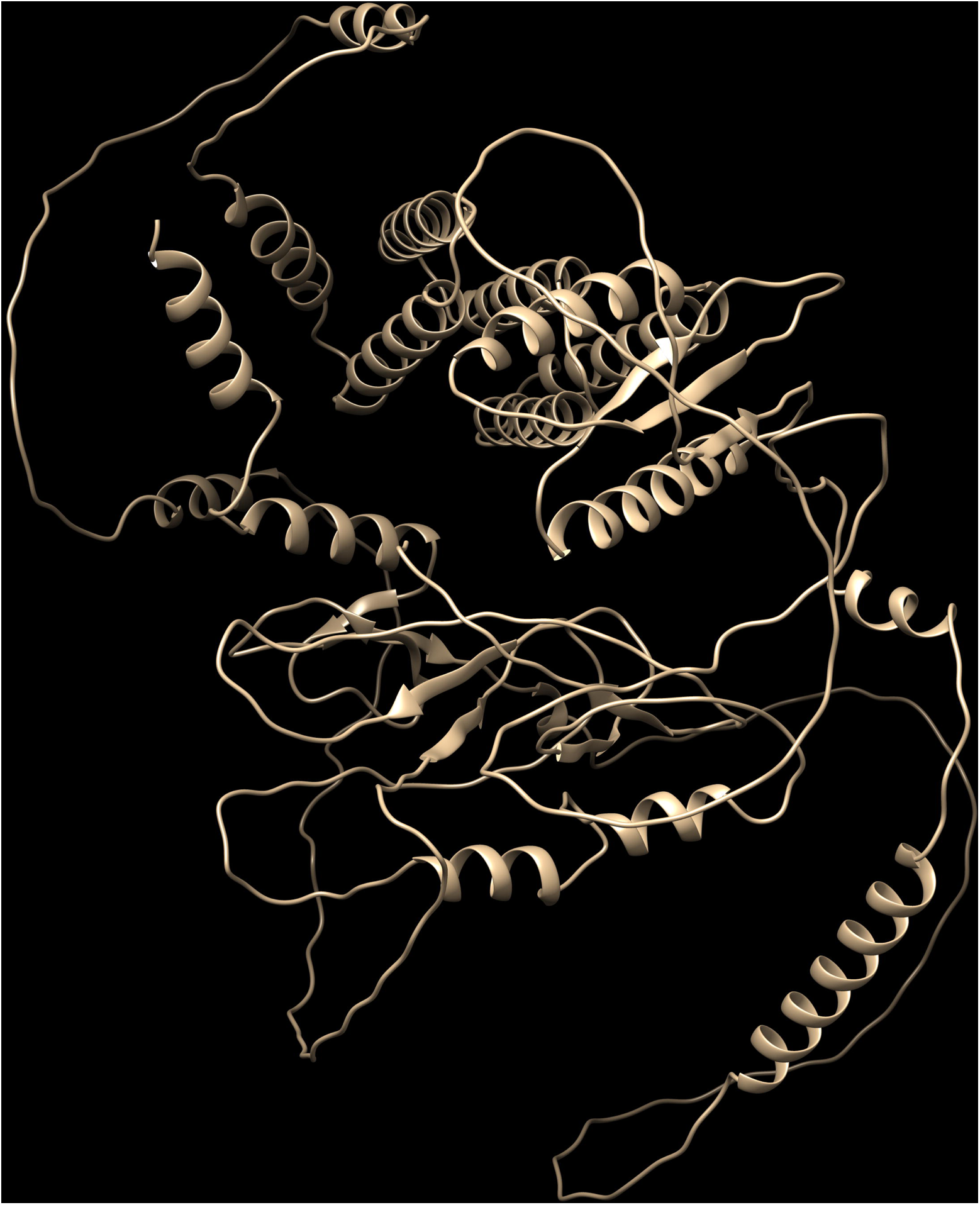
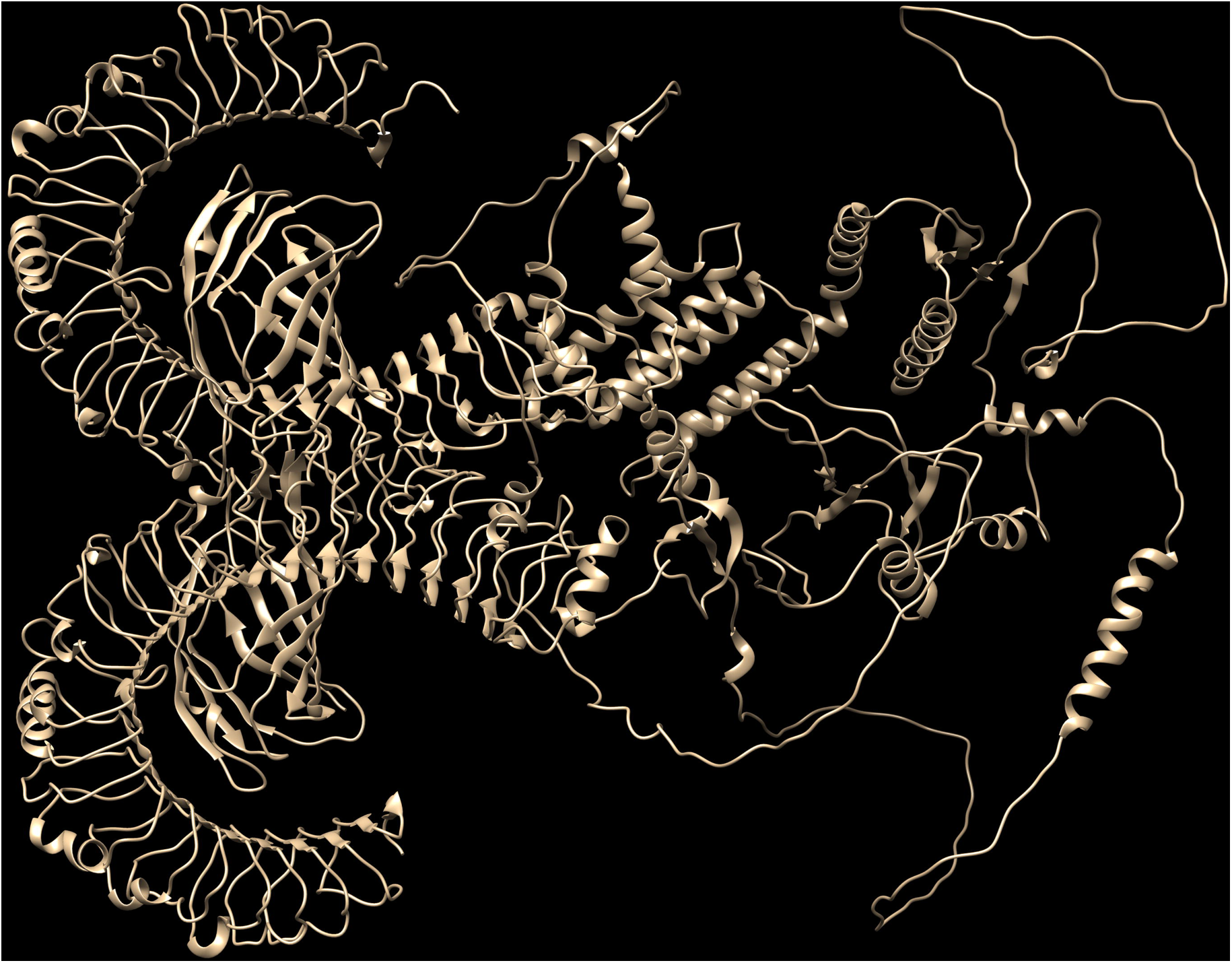
(A): The final sequence of the designed multi-epitope vaccine and adjuvants. The EAAAK linker is present between the adjuvant and B-cell epitopes with KK linkers. CTL epitopes have AAY linkers, while HTL epitopes have GPGPG linkers. 1 (B) Diagrammatic representation of the final vaccine.

### Prediction of the vaccine peptide’s antigenicity, allergenicity and toxicity

The VaxiJen 2.0 server was used to predict the antigenicity of the final sequence; the results were 0.8977 using a bacteria model and 0.5195 using the parasite model at a threshold of 0.4. According to the results obtained, the produced sequences, both with and without adjuvant, are antigenic. The vaccine sequence was also predicted to be non-allergenic using the AllerTOP v.2 server, with and without the chosen adjuvant. The CSM Toxin server was utilized to predict the toxicity. The complete peptide sequence was found to be non-toxic.

### Physicochemical properties of the vaccine peptide

The final protein’s molecular weight (MW) was 80.77415 kDa, with a total of 751 amino acids. The theoretical pI was 9.75, and it was determined that the half-life was 30 hours in mammalian reticulocytes, >20 hours in yeast, and >10 hours in E. coli in vivo. It was predicted that the aliphatic index was estimated to be 60. 03, suggesting thermostability. The grand average of hydropathicity (GRAVY) was predicted to be -0.758. The negative value shows that the protein interacts with water molecules and is hydrophilic. An instability index of 33.04 was obtained, indicating that the protein is stable.

### Solubility prediction of the vaccine peptide

With a solubility score of 0.641 at a threshold of 1.6 and a Kyle-Doolittle hydropathy value of 1.98, it was projected that the protein would be soluble upon expression.

### Prediction of secondary structures

The PSIPRED server was used to predict the protein’s secondary structure. The final structure of the chimeric peptide was predicted to comprise 25% alpha helix, 10% beta strand, and 65% coil. The results suggested a flexible multi-epitope vaccine with good immune recognition. Figures 2 (A) and 2 (B) represent the PSIPRED pictorial prediction of secondary structure.

**Figure 2:**
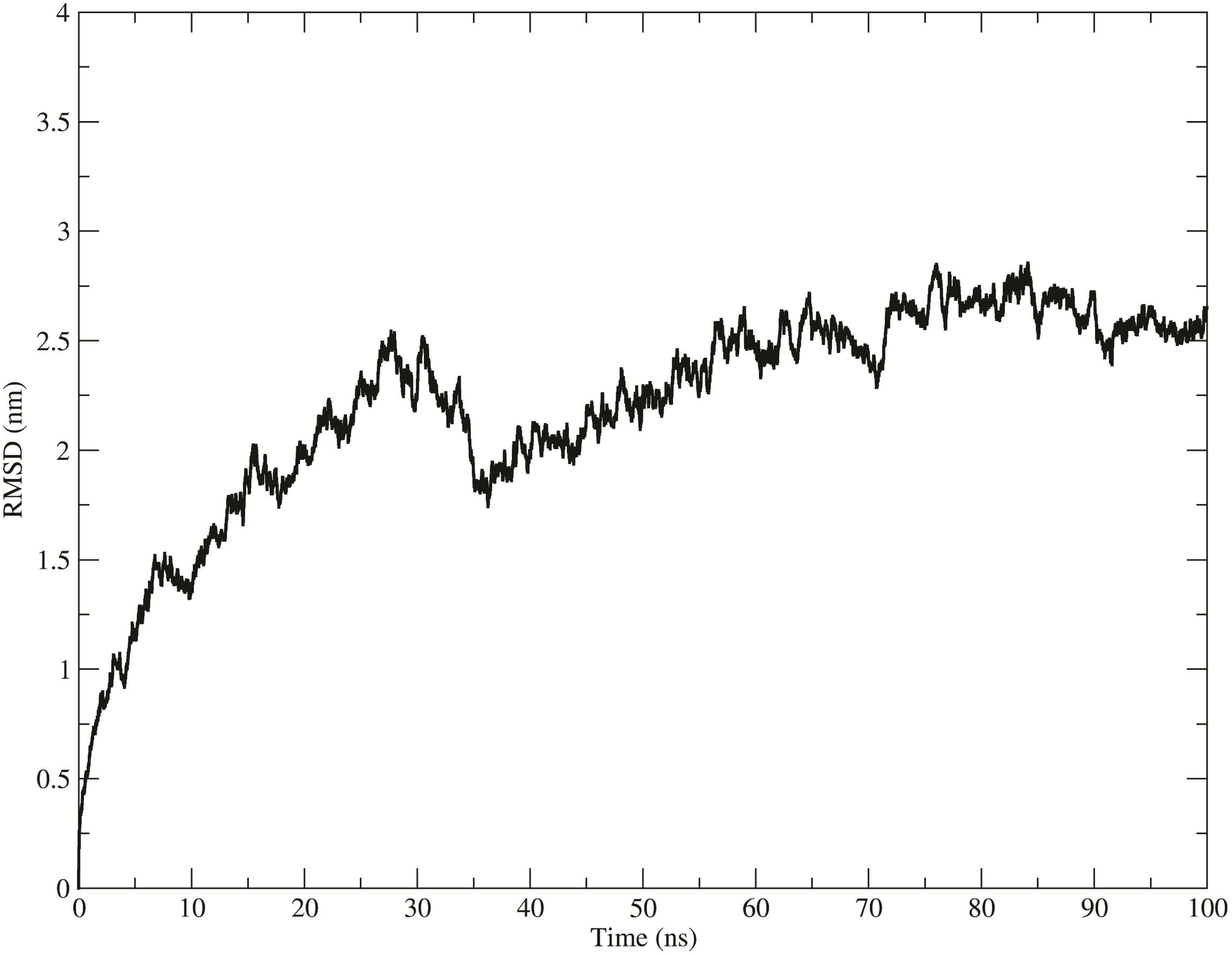
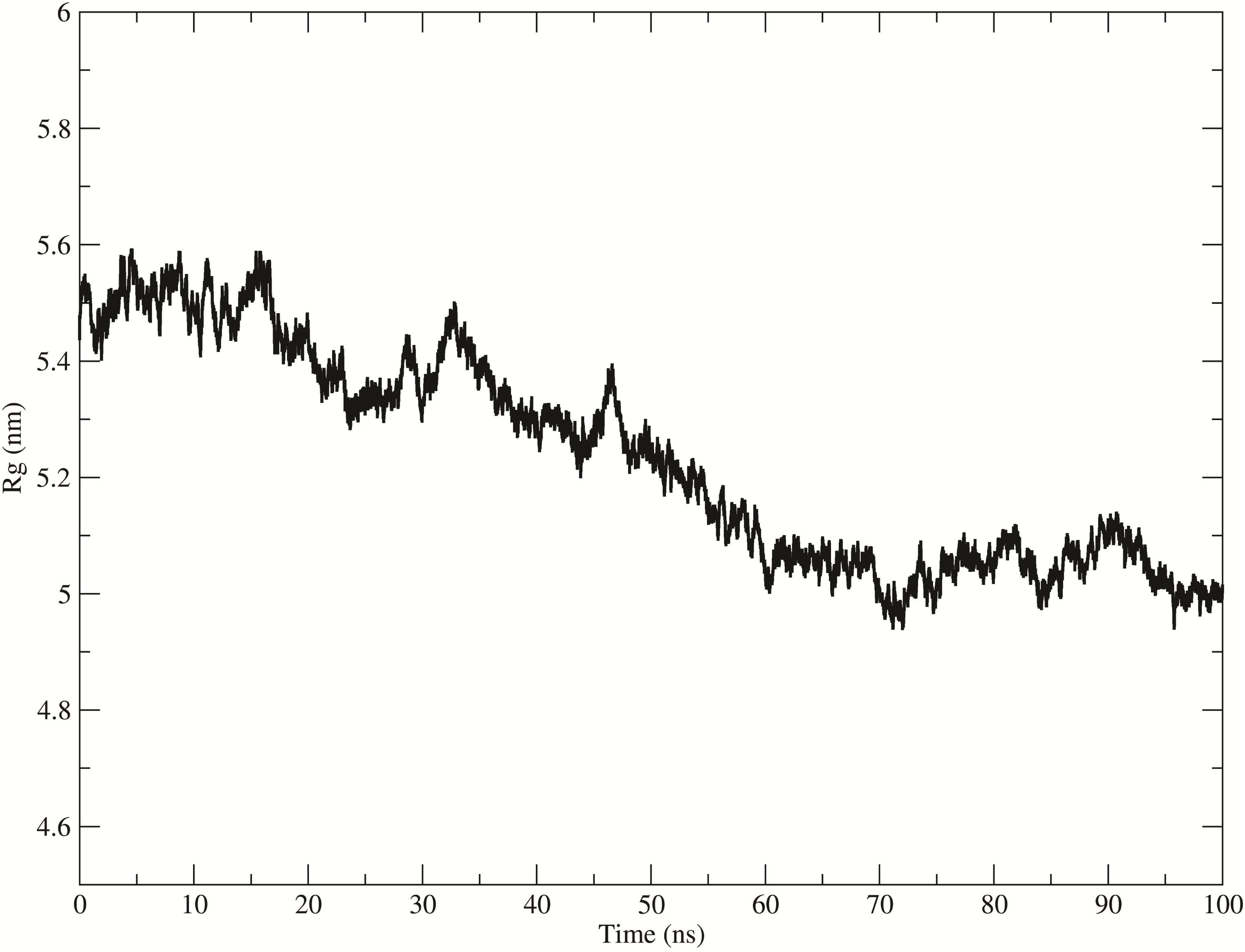
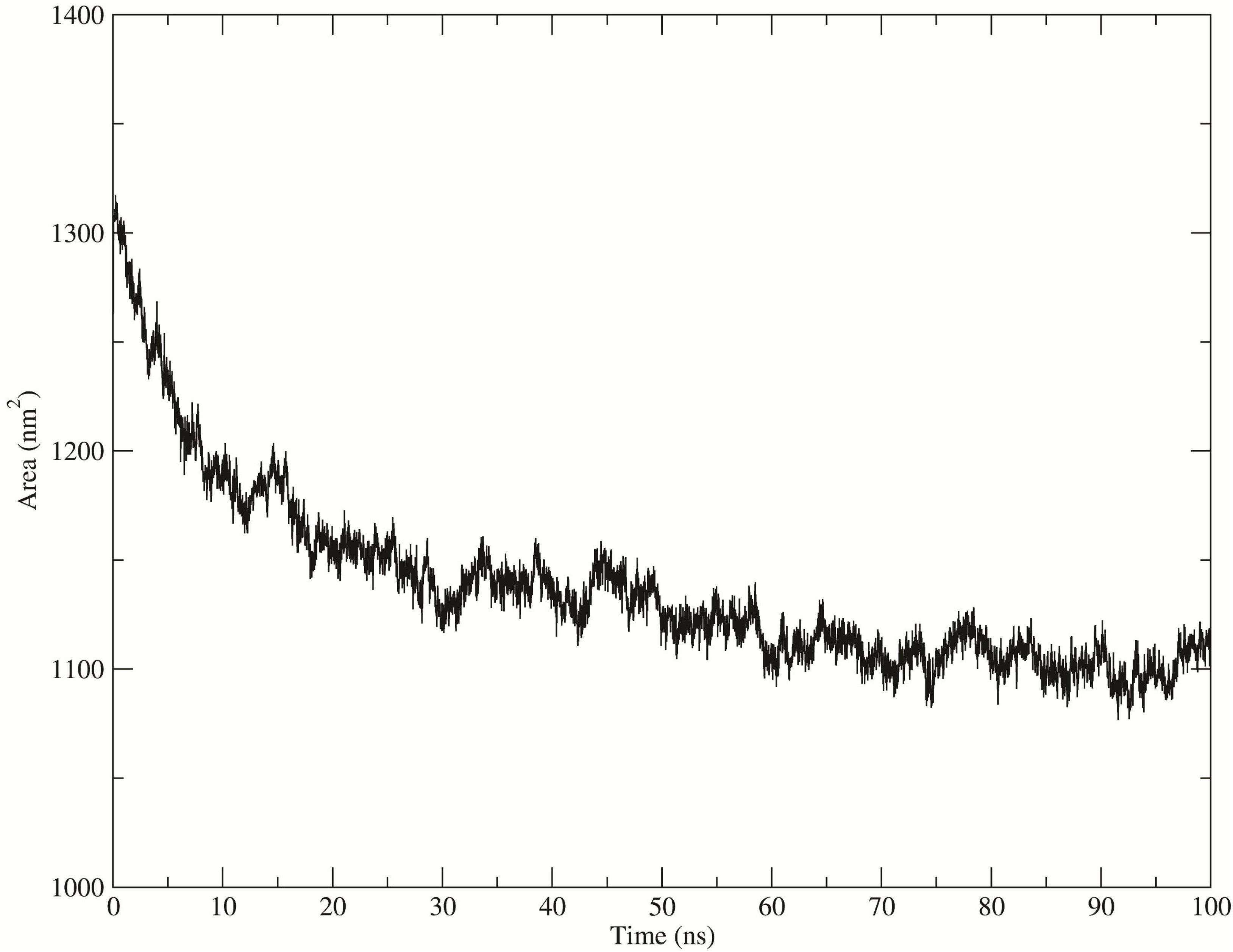
PSIPRED results of the predicted secondary structure of the multi-epitope vaccine construct. (A) and (B) Shows the secondary structure for all amino acids with the strands in yellow, helices in pink and coils in grey. All the selected epitopes and the linkers are included. ‘Conf’ shows the confidence of prediction for the secondary structures, supported by the most probable assignment of every residue which is given by ‘pred’. High prediction confidence is seen in most secondary structures and is more notable in coils.

### Prediction, refinement, and validation of tertiary structure

The tertiary structure of the proteins was predicted and validated using the AlphaFold software. The Ramachandran plot was created with the Molprobity server. For model refinement, the GalaxyRefine server was used; five vaccination models were produced by refining the original “crude” model. Model 3 was determined to be the most suitable based on the model quality scores for all revised models. This was hence chosen as the final vaccine model to be considered for further analysis. The modelled protein’s Ramachandran plot study showed that 99% of its residues are located in preferred areas. This aligns with the projected 97% score from the GalaxyRefine analysis. Furthermore, only 1% of the residues were anticipated to be in disallowed locations, while all others were expected to be in allowed regions. The chosen model no. 3 from the GalaxyRefine server is shown in Figure 3; the TLR 4+ Model 3 structure is depicted in Figure 4, and the Ramachandran plot is shown in Figure 5.

**Figure 3:**
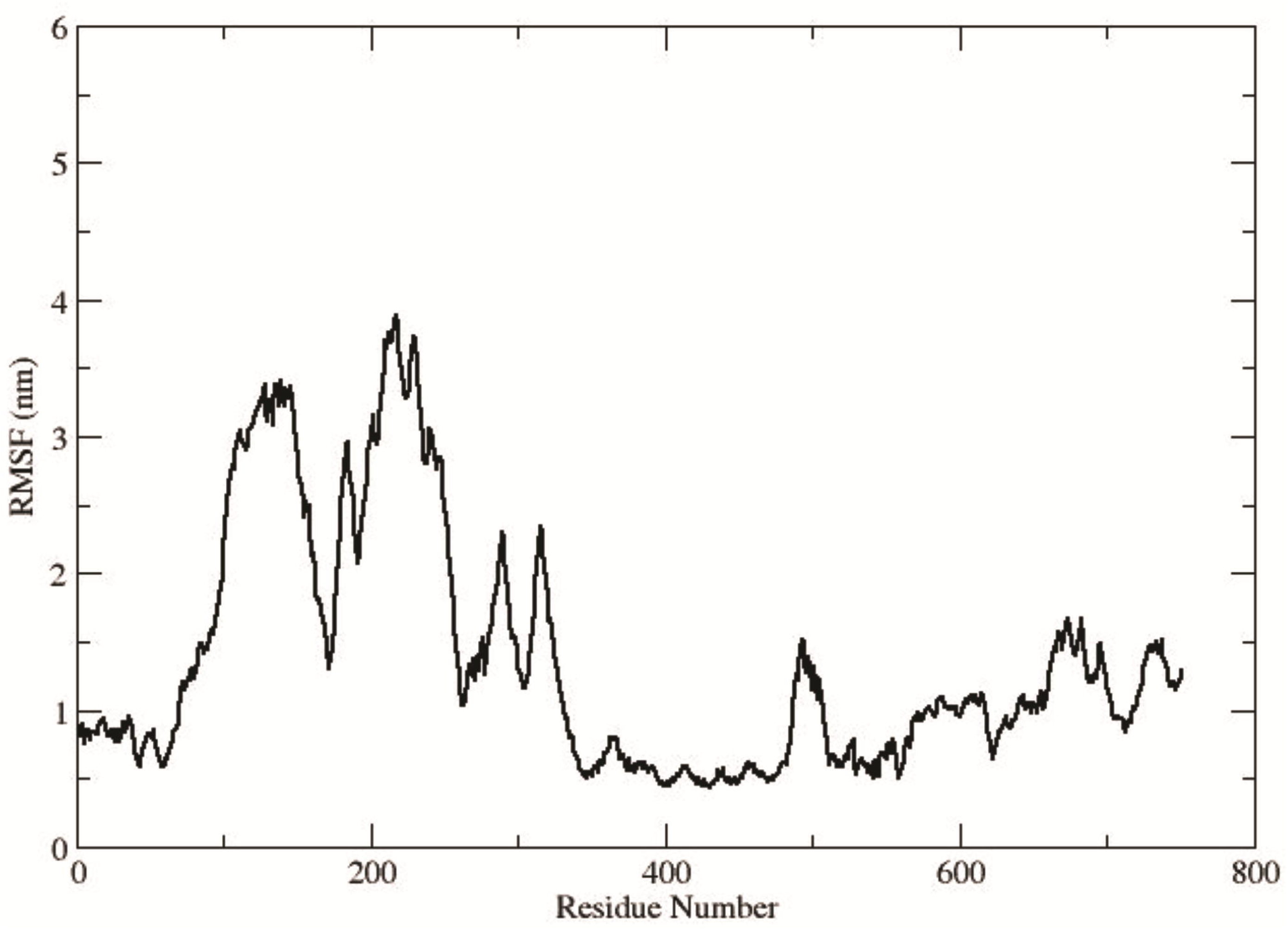
Tertiary structure of the multi-epitope vaccine after modelling and refinement. The selected model was obtained using GalaxyRefine. The top ranking model was selected and was viewed using Chimera structural viewer.

**Figure 4:**
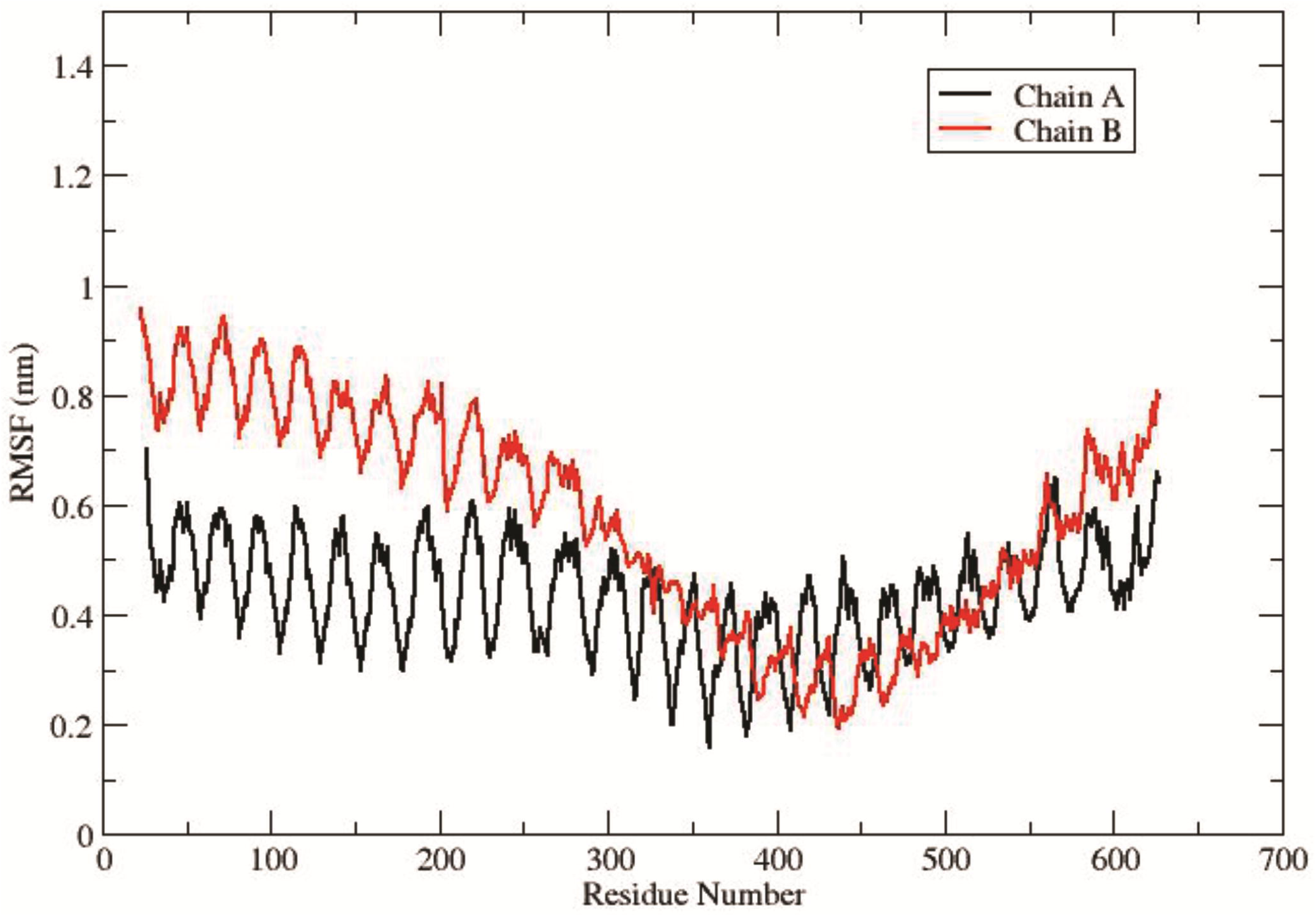
Tertiary structure of TLR 4 and vaccine model of the multi-epitope vaccine after modelling and refinement. The selected model was obtained using GalaxyRefine, and was viewed using Chimera structural viewer.

**Figure 5:**
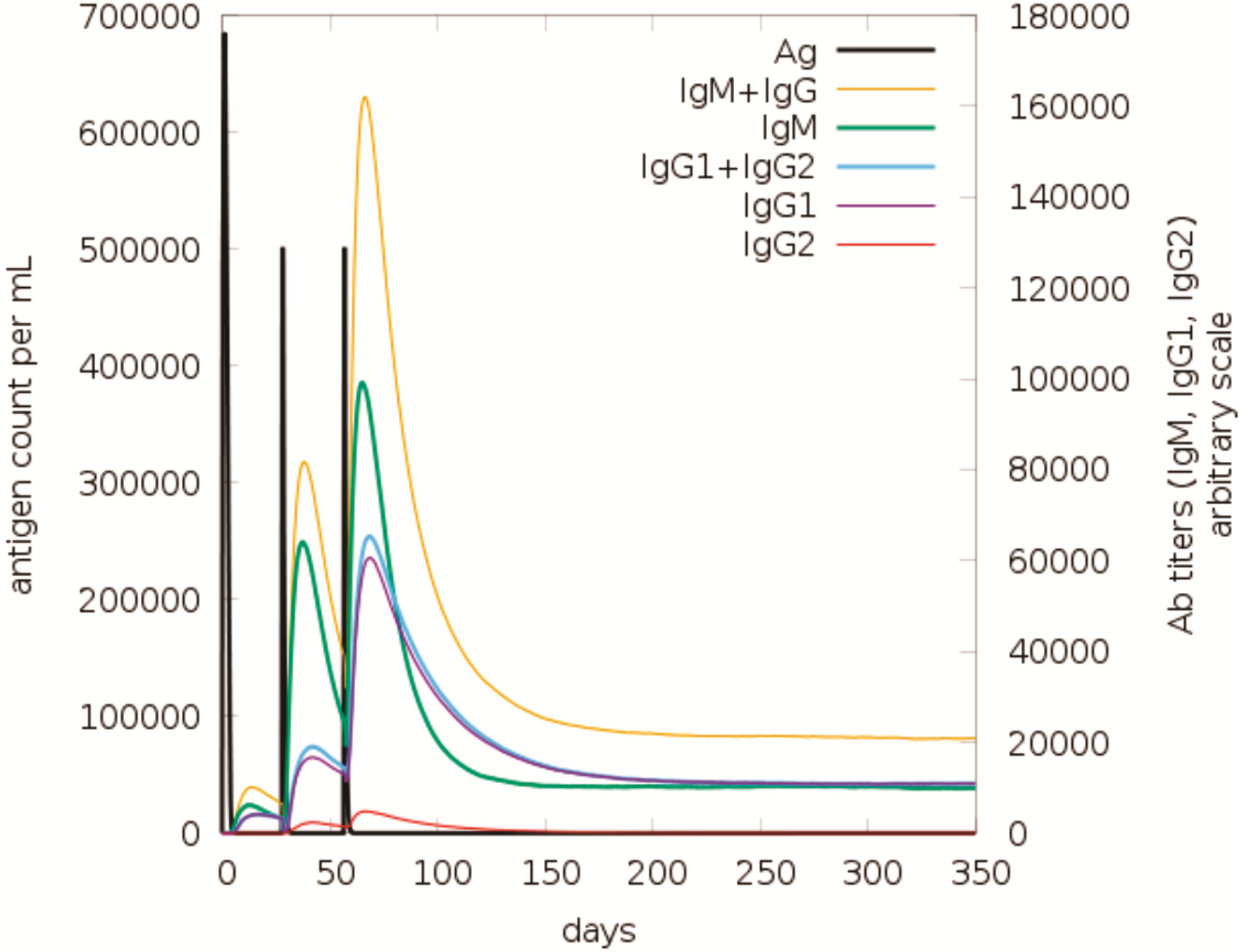
Shows validation of multi-epitope vaccine tertiary structure by Ramachandran plot generated using Molprobity software, showing 99% of the residues in preferred areas and only 1% in disallowed regions.

### Molecular dynamic (md) simulations

The MD simulation trajectory data provide valuable insights into the binding mode and stability of the TLR-vaccine complex Figure 4. The analysis begins with the evaluation of the root mean square deviation (RMSD) of the backbone atoms, which reflects the overall stability of the complex over time. The RMSD plot, as shown in Figure 6 (A), indicates that the complex remains stable throughout the simulation after 70ns, with fluctuations remaining below 3 nm. This suggests that the TLR receptor and the vaccine construct maintain a consistent and stable interaction throughout the 100 ns simulation period.

**Figure 6:**
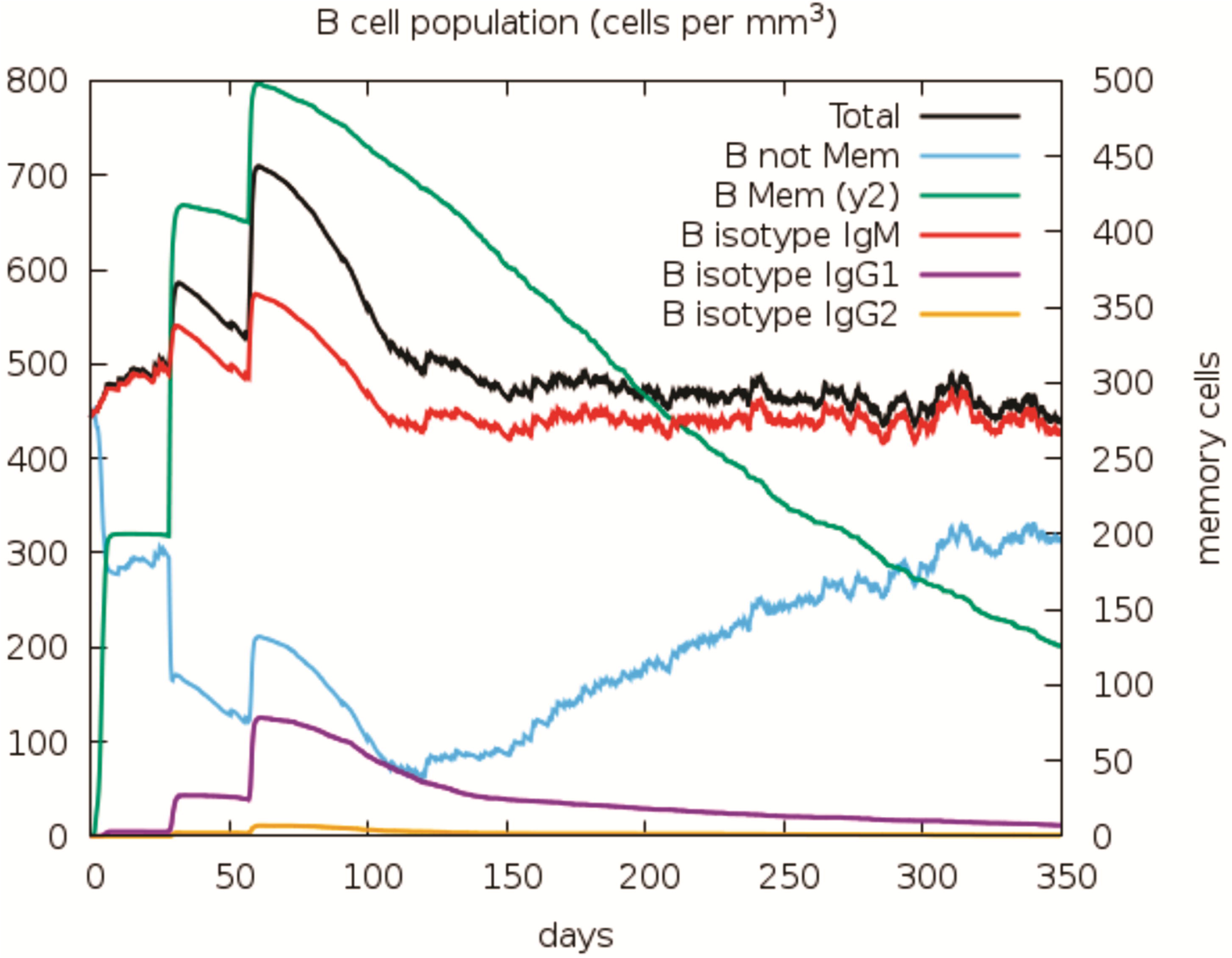
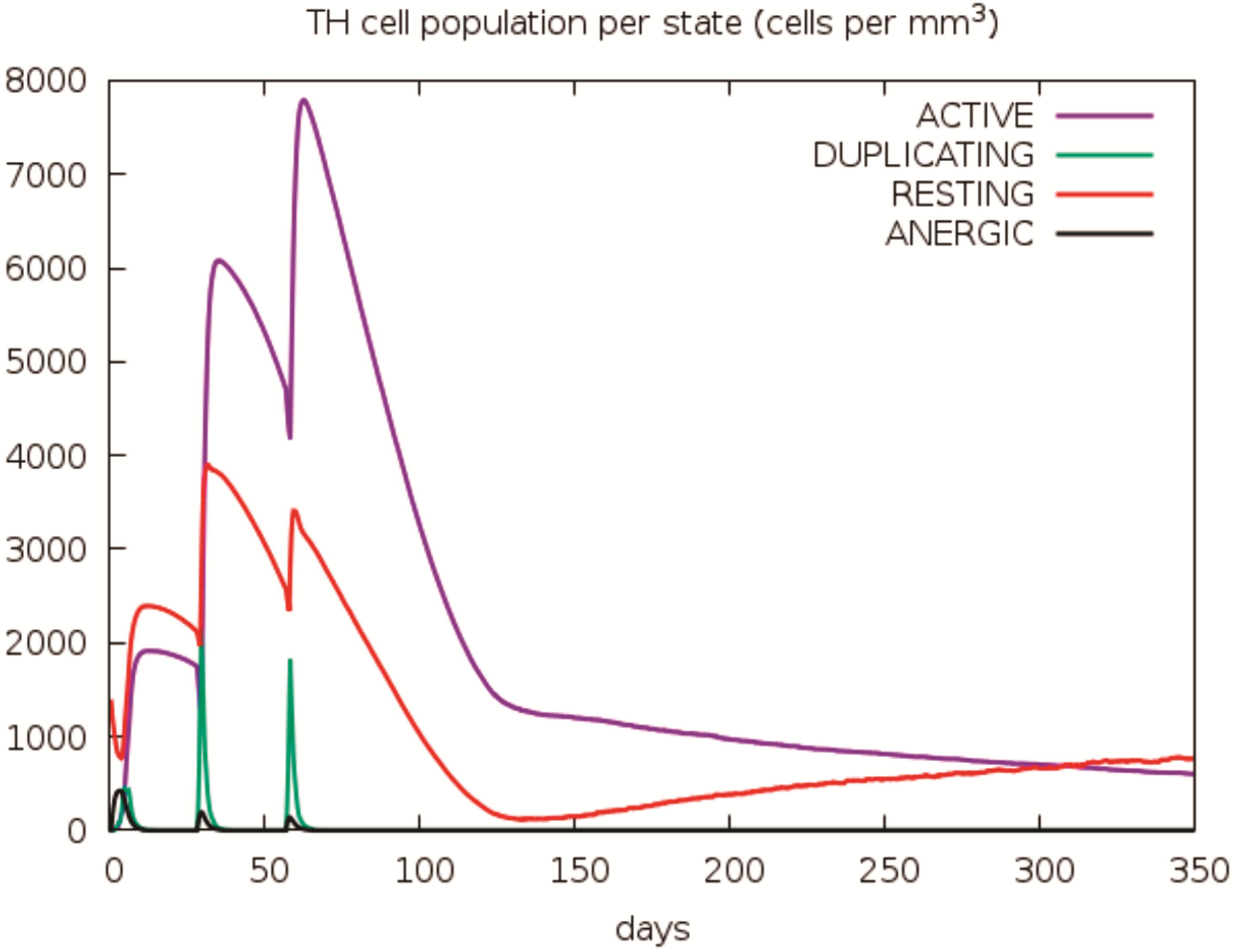
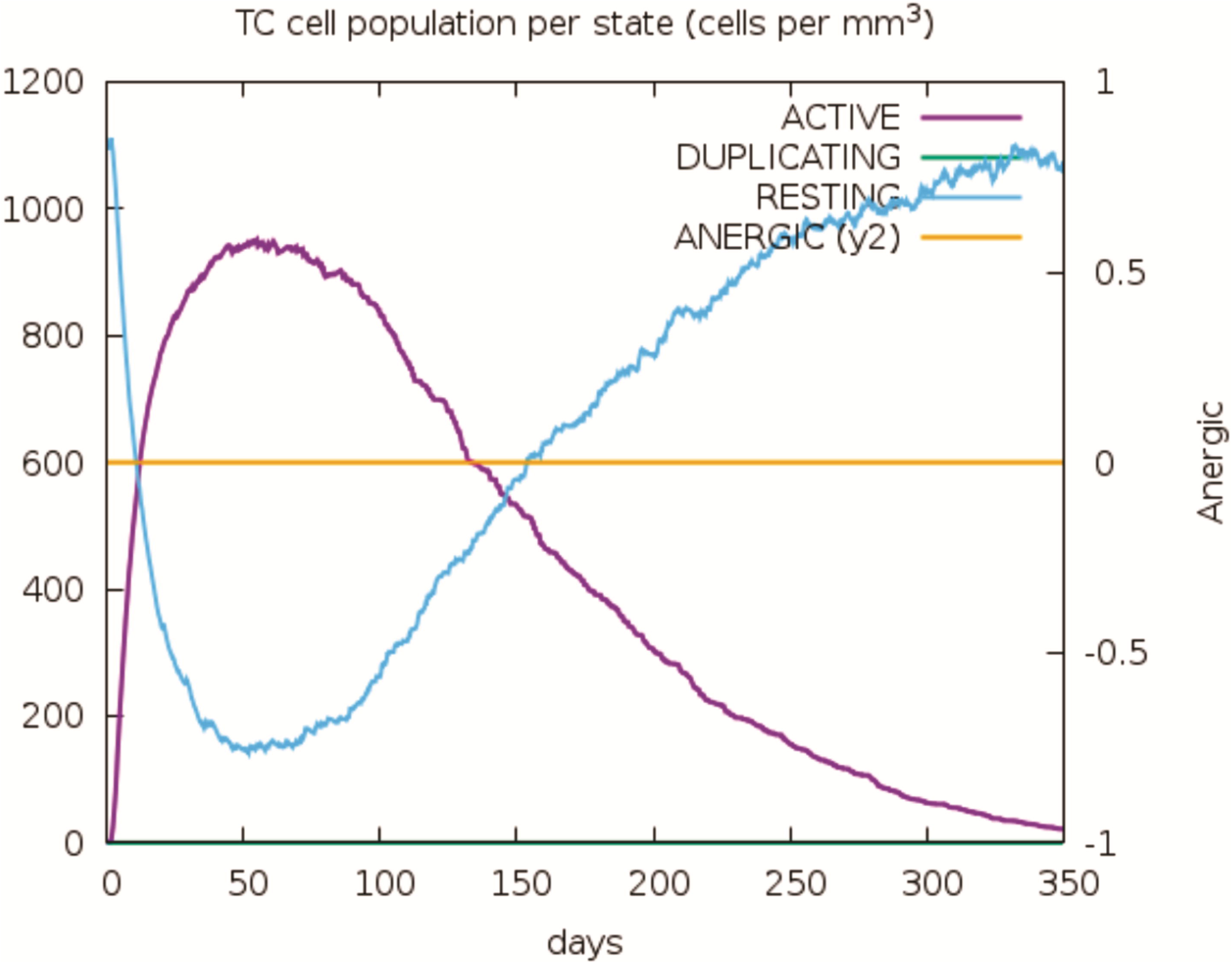
Root mean square deviation (RMSD) and radius of gyration (Rg) plots for the TLR-vaccine complex. (A) The RMSD plot indicates that the TLR receptor and vaccine construct form a stable complex throughout the simulation. Similarly, (B) the Rg plot demonstrates the compactness and stability of the complex, (C), SASA plot reflects the compactness of the vaccine and TLR complex, further supporting its structural integrity during the MD simulation.

Next, the compactness of the complex was assessed using the radius of gyration (Rg), shown in Figure 6 (B). The Rg values, which measure the compactness of the protein complex, demonstrated a stable state with a gradual decrease in values from 5.6 nm to 5.0 nm over the simulation. This reduction in Rg indicates that the complex adopts a more compact conformation as the simulation progresses, suggesting that the TLR receptor and the vaccine construct come closer together, likely due to favorable interactions stabilizing the structure.

The solvent-accessible surface area (SASA) analysis further supports the stability and compactness of the complex. Initially, the SASA values were high (1300 nm²), indicating an exposed and loosely packed structure. However, after 50 ns, the SASA values stabilized at around 1100 nm², signifying a tighter and more compact packing of the complex as indicated in Figure 6 (C). This reduction in surface area suggests that the complex becomes more stabilized as hydrophobic regions are buried, reducing exposure to the solvent.

Root mean square fluctuations (RMSF) were also calculated for TLR chains A and B, as well as for the vaccine construct, to assess the flexibility of specific regions within the complex. The RMSF plot highlights the movement of Cα atoms during the simulation. Regions exhibiting higher RMSF values correspond to flexible regions, while lower RMSF values indicate more constrained regions. Notably, residues within the 350-800 region of the vaccine construct displayed minimal fluctuations, suggesting that these residues are critical for binding to the TLR receptor and contribute to the stability of the interaction, demonstrated in Figure 7 (A). Similarly, TLR chain A and B residues in the 300-600 region exhibited lower fluctuations, reinforcing their role in stabilizing the complex by maintaining a stable interface with the vaccine construct, shown in Figure 7 (B).

**Figure 7:**
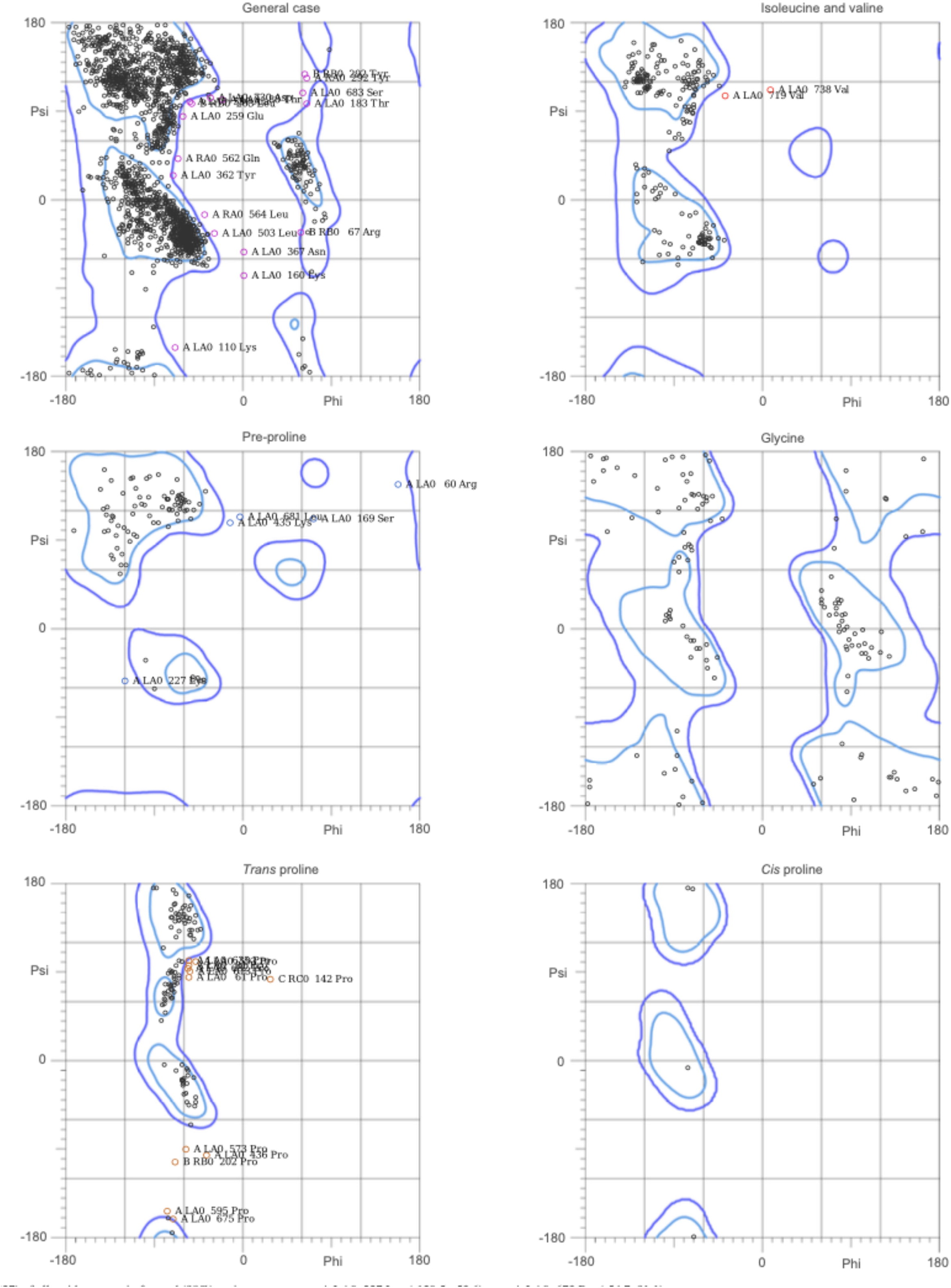
Root mean square fluctuations (RMSF) of the TLR-vaccine complex. (A) illustrates the RMSF of the vaccine construct, highlighting regions of flexibility and stability. (B) depicts the figure of the TLR receptor, with chain A shown in black and chain B in red, over the 100 ns simulation. These plots reveal the dynamic behaviour of the complex, where flexible and stable regions contribute to binding and overall structural integrity.

Overall, the MD simulation trajectory data indicate that the TLR-vaccine complex remains stable, compact, and tightly packed throughout the simulation. The minimal fluctuations observed in the key regions of both the vaccine construct and the TLR receptor suggest a strong and stable binding interaction, providing important insights into the molecular basis of the TLR-vaccine binding and its potential for further optimization in vaccine design.

### Binding energy calculations

The binding energy calculations using the MM-GBSA method offered detailed insights into the energetic factors governing the interaction between the TLR receptor and the vaccine construct. These calculations were based on the equilibrated portion of the molecular dynamics trajectory, starting from 50 ns. The total binding energy was calculated to be - 245.83 kcal/mol, indicating a strong and stable interaction between the two components. The Van der Waals energy contribution (-403.51 kcal/mol) demonstrated favorable dispersion forces arising from non-covalent atomic interactions, suggesting good steric complementarity and a stable hydrophobic interface that helps maintain the structural integrity of the complex. Electrostatic interactions contributed significantly to the binding affinity, with a large negative energy value (-9959.94 kcal/mol) reflecting strong charge-charge attractions critical for stabilizing the TLR-vaccine interface. Conversely, the Generalized Born (GB) energy contribution (10175.03 kcal/mol) represented the polar solvation energy and highlighted a substantial desolvation penalty when polar regions of the molecules were buried upon binding, partially offsetting the electrostatic interactions. The Solvent-Accessible Surface Area (SURF) energy contribution (-57.41 kcal/mol) indicated favourable hydrophobic effects, as water molecules were displaced during the interaction of hydrophobic surfaces between the receptor and the vaccine construct. The gas-phase energy (GAS, -10363.44 kcal/mol), which combines van der Waals and electrostatic contributions in the absence of solvent, revealed highly favourable direct interactions between the components. Meanwhile, the total solvation energy (10117.62 kcal/mol) accounted for both polar and nonpolar solvation costs, reflecting the energetic penalty associated with desolvating charged and polar regions.

The net binding energy of -245.83 kcal/mol underscores a balance between favourable gas-phase interactions and the unfavourable solvation penalty, with electrostatics playing a dominant role in the binding interface. Additionally, favourable van der Waals and hydrophobic contributions further enhanced the complex’s stability. These results provide a comprehensive understanding of the energetics underlying the interaction and offer a foundation for optimizing vaccine design by targeting key residues involved in electrostatic and van der Waals interactions. Overall, the MM-GBSA analysis confirms that the TLR receptor and the vaccine construct form a stable and energetically favourable complex.

### Predictive analytics of immune simulations

The C-ImmSim server was utilized to evaluate the vaccine protein’s immune response and immunogenicity. The strength and quality of these two factors are in determining vaccination efficacy. A vaccine with a high level of immunogenicity has a greater chance of preventing the targeted disease. The server predicted how vaccination candidates interact with immune receptors, enhancing the responses of both the humoral and cellular immune systems. Figure 8 illustrates the C-ImmSim presentation of the *in-silico* immune simulation featuring the chimeric peptide.

**Figure 8:**
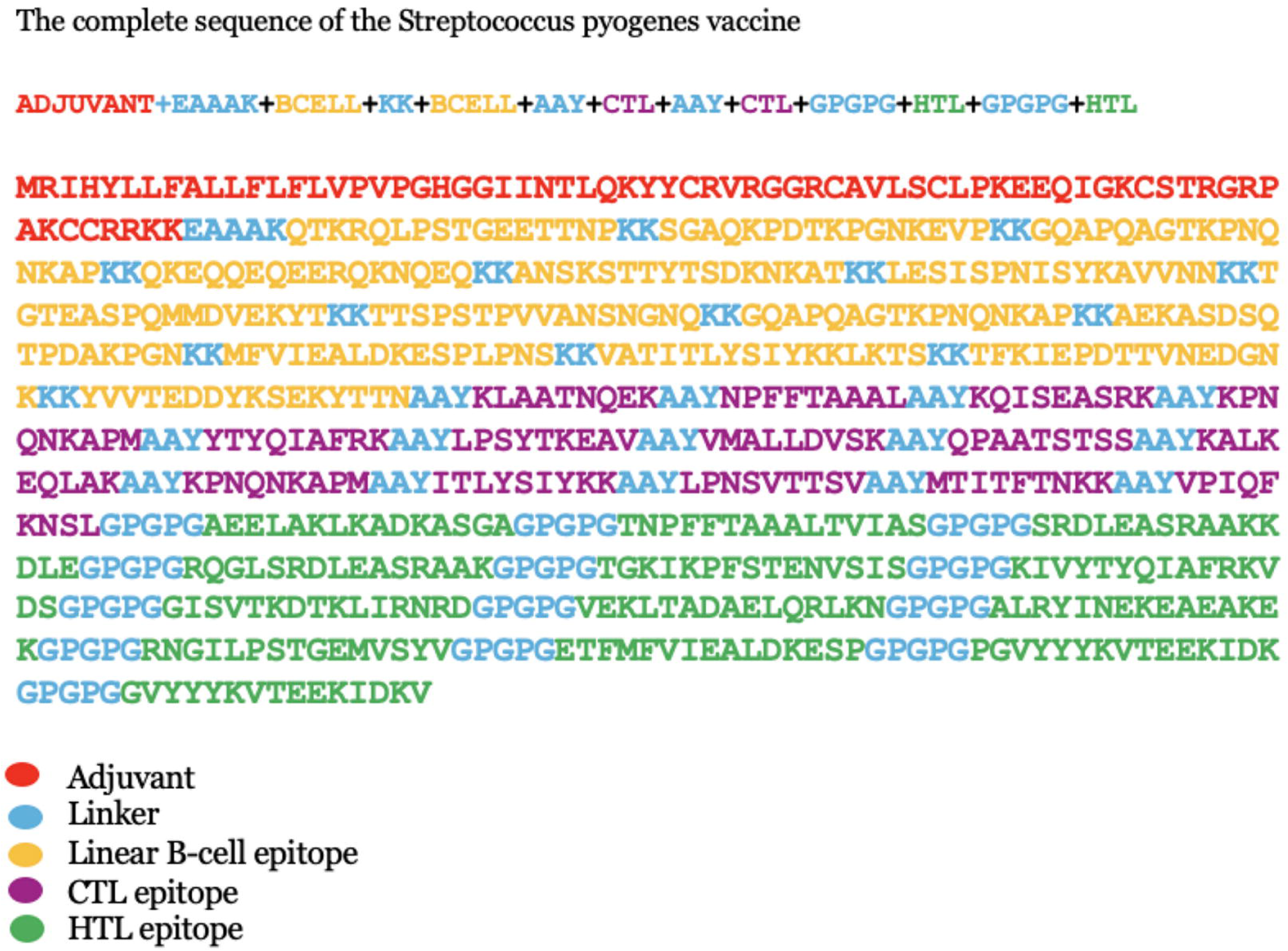
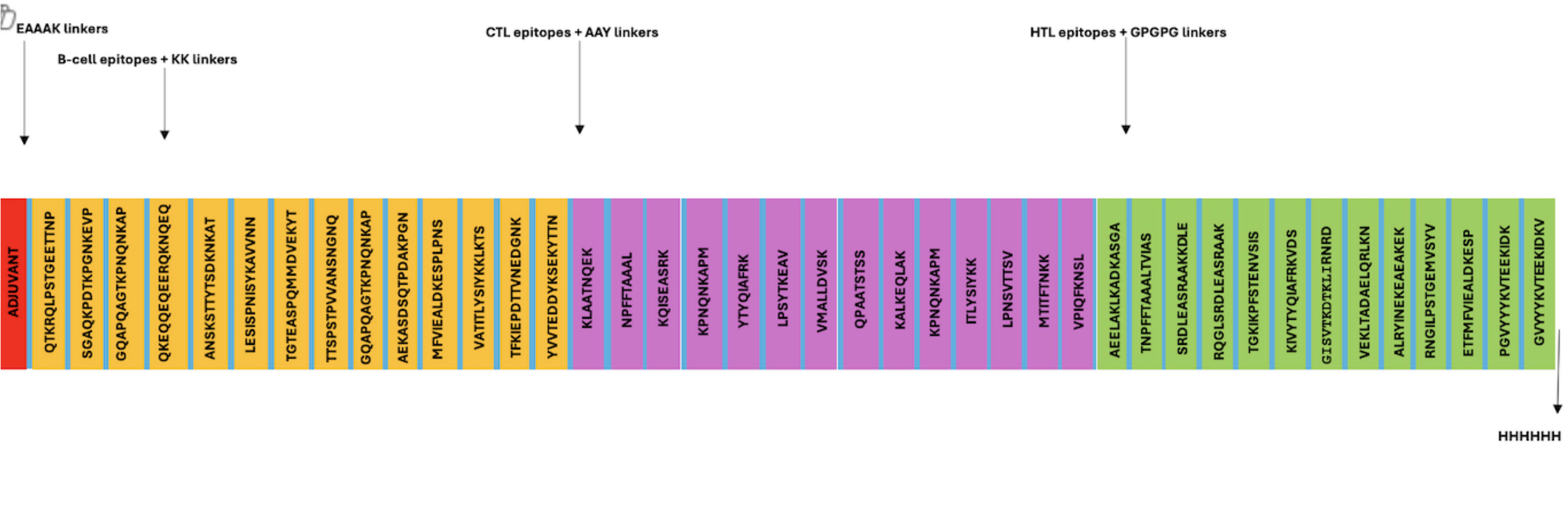
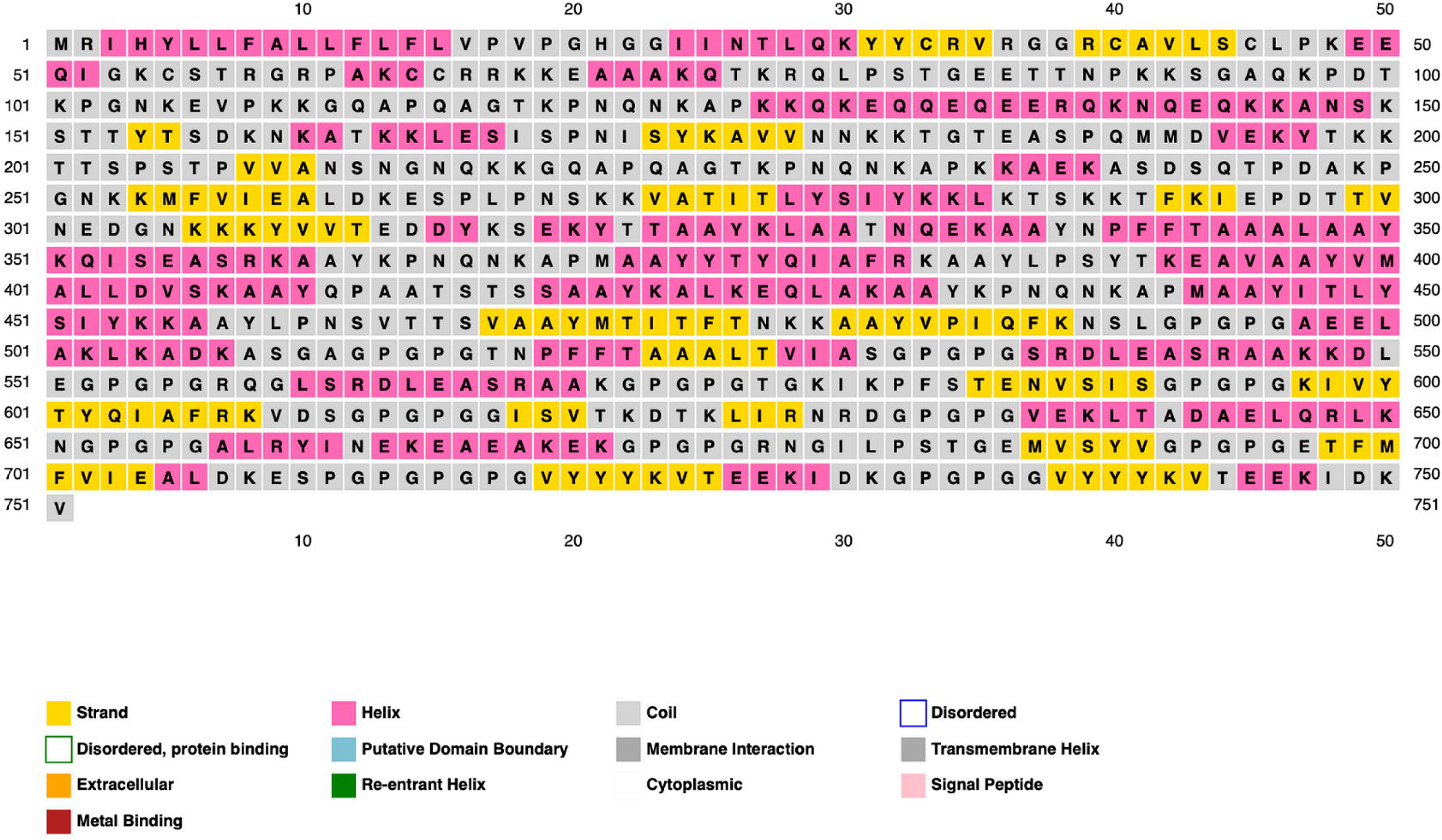
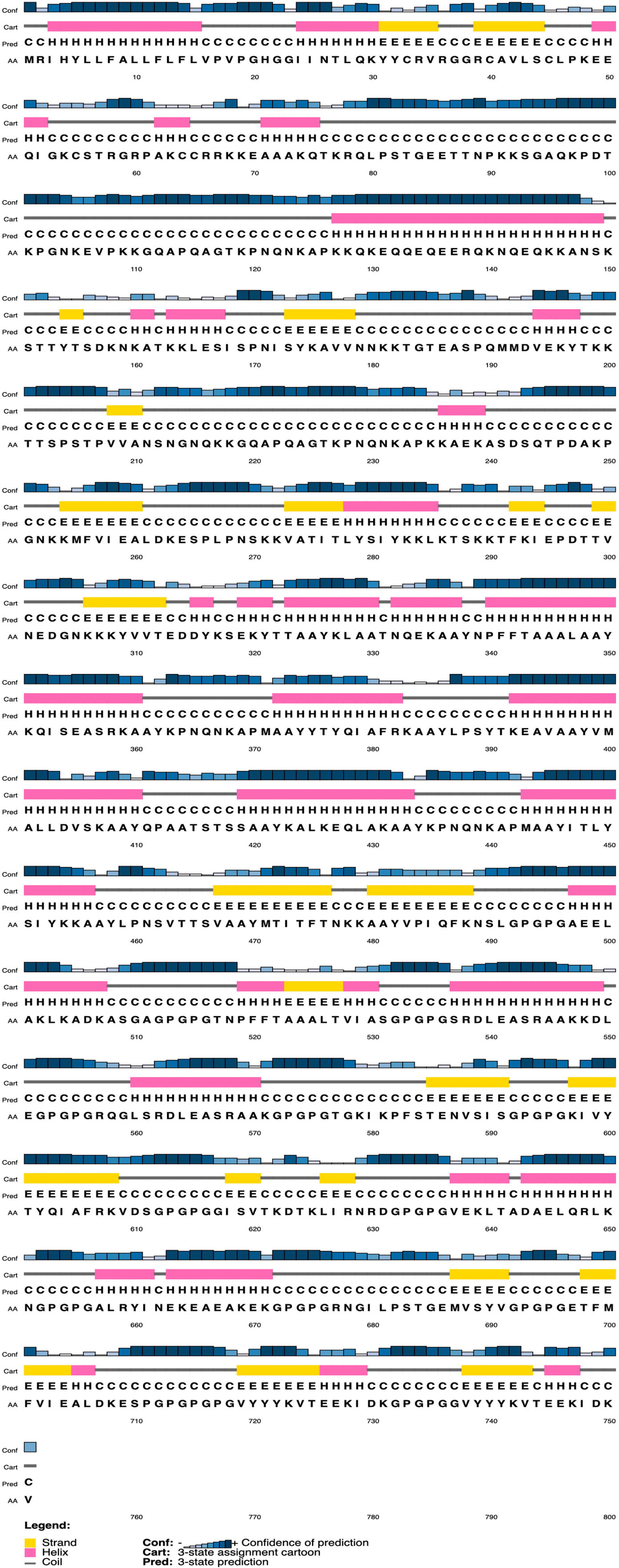
C-ImmSim immune simulation of the chimeric peptide. (A) Synthesis of immunoglobulins due to antigen injection which is represented by vertical black lines. (B) Changes in the population of B-cell after the three injections. (C) Changes in TH cell population after injections. (D) Change in TC cell population after injections.

Black vertical lines indicate the levels of antigen injections. IgM (green), IgG1 and IgG2 (blue) showed increased and higher activity immediately after each injection. Increased duration of response by IgG indicates effective clearance of antigen, shown in Figure 8 (A). The homologous IgM B cells, and memory B cell levels significantly increased after each injection, but memory B cells activity slowly declined after the third, suggesting the formation of long-term memory. Homologous IgG1 B cells also increased each time, but at a lower rate, while homologous IgG2 B cell showed little change, illustrated in Figure 8 (B). The active and resting T helper cell levels increased after each injection, showing greater activity each time, shown in Figure 8 (C). The resting T-cell levels decreased after the first two injections, but increased after the third. In contrast, the active T-cell showed higher activity after the first two injections, but lowered after the third, seen in Figure 8 (D). The anergic state signifies the T-cells’ tolerance to the antigen caused by repeated exposures. In contrast, the resting state denotes inactive cells that were not exposed to the antigen.

### Codon Optimization and Reverse Translation

The Java Codon Adaptation Tool (JCat) was used to optimize codon usage for maximum protein expression in the vaccine design within E. coli (strain K12). The optimized codon sequence consists of 2,253 nucleotides. This improved nucleotide sequence has a Codon Adaptation Index (CAI) of 0.98, with a GC content of 49.3%, compared to the GC content of Escherichia coli (strain K12), which is 50.7%. The recommended GC content range is between 30% and 70%. Overall, the results indicate that the vaccine is highly optimized for expression in the host E. coli.

## DISCUSSION

*S. pyogenes* is among the top ten fatal human infectious diseases. It manifests in a wide range of clinical symptoms due to its ability as an opportunistic pathogen (57). The *emm* gene coding the M protein is highly variable, resulting in more than 250 *emm* types being reported worldwide. High-income regions tend to have a lower strain diversity, in contrast to low-income regions indicating the relationship between social factors, epidemiological diversity and occurrence of the disease (58). In addition, the bacterium’s ability of molecular mimicry raises concerns of autoimmunity and high reactogenicity post administration make it challenging to create a vaccine (59). Multi-epitope based vaccines are more effective due to the presence of multiple MHC-restricted epitopes which activate T-cells while also inducing greater immune response by incorporating B-cell, CTL, and HTL epitopes. Addition of adjuvants and targeted antigens strengthens immunogenicity and long-term recognition (60). This study used proteins of *S. pyogenes* and employed numerous immune-informatics methodologies to create a potential MEBV vaccine.

We began by sorting the various sequences of the M protein. Five *emm* proteins were chosen based on prevalence and overall coverage. Four components of the immune system were then selected for use as epitopes: B-cell, cytotoxic T-cell, helper T-cell and IFN- γ. ABCPred server was used to predict B-cell epitopes, while CTL and HTL epitopes were predicted using NetCTL-1.2, and NetMHCII 2.2 respectively. Possible IFN- γ epitopes were selected with IFNepitope server. The optimal sequences were used to construct the vaccine candidate in combination with the adjuvant Beta-defensin to increase its immunogenicity. Linkers KK, AAY and GPGPG were added between the epitopes and adjuvants appropriately. Adjuvants stimulate both the innate and adaptive immune response, while linkers are vital for providing stability and cross-domain interaction (61).

VaxiJen v2.0 server was utilized to predict the antigenic reactivity, AllerTOP v2.0 for allergenicity, and toxicity was tested using CSM-Toxin service. The construct was found to stimulate antigenicity, was non –allergenic and non-toxic. Molecular dynamic simulations were then used to study the behaviour of the construct in a real biological system by assessing the structural stability and physical movement under different conditions (62). The results indicate the potential of the vaccine to engage TLR receptors to form a stable complex. The ability of the vaccine construct to stimulate the immune system was conducted by the C-ImmSimm server (63). Simulated results show effective Ig synthesis and activation of T-helper and T-cytotoxic cells. Finally, JCat was to optimize the codon sequence to ensure maximum protein expression.

This study developed a potentially effective vaccine for *S.pyogenes* as it is a multi-epitope vaccine comprised of B-cell, HTL, CTL and IFN- γ epitopes. The construct is capable of stimulating a strong and long-term immune response, while demonstrating considerable antigenicity, but displaying no toxicity or allergenicity.

## CONCLUSION

Public awareness of *S. pyogenes* has recently increased due to its re-emergence as a source of serious human infections in the USA, Europe, and other countries, as extensively documented over the past few decades. The absence of an approved *S. pyogenes* vaccine is a significant concern and emphasizes the need to enhance global surveillance of this pathogen. This study presents a potential vaccine peptide coding for several B-cell and T-cell (HTL and CTL) epitopes designed using immunoinformatics methods. The vaccine peptide may provide both preventive and therapeutic benefits since the proteins with these epitopes are expressed during the pathogen’s infectious stages. This chimeric vaccine peptide could serve as an additional strategy to eradicate *S. pyogenes*. New investments, advancements in developing current vaccination candidates, and the introduction of new candidates have made progress in this area. However, the vaccine pipeline remains limited, and all aspects of vaccine development must be accelerated to meet the strong demand for an effective vaccine.

## AUTHOR CONTRIBUTIONS

Concept and design**: RKP;** Data acquisition, analysis, interpretation, manuscript drafting and diagrammatic representation: **SH**. Manuscript revision: SH, SMR, and RKP; Project supervision: RKP

## ACKNOWLEDGEMENTS

We express our appreciation to Dr. Bajarang Kumbhar for his contribution and guidance in the molecular dynamic simulation.

## CONFLICT OF INTEREST

The authors declare that they have no conflict of interest.

## FUNDING

The authors received no financial support for the research and publication of this article.

## Notes

### Competing Interest Statement

The authors have declared no competing interest.

